# Stemness activity underlying whole brain regeneration in a basal chordate

**DOI:** 10.1101/2021.10.24.465595

**Authors:** Tal Gordon, Tal Zaquin, Mark Alec Kowarsky, Yotam Voskoboynik, Noam Hendin, Omri Wurtzel, Federico Caicci, Lucia Manni, Ayelet Voskoboynik, Noa Shenkar

## Abstract

Central nervous system (CNS) regeneration extent is highly diverse across the metazoans, with adult mammals demonstrating limited ability^1,2^. Understanding how neurons regenerate following injury remains a central challenge in regenerative medicine. Although conserved pathways associated with neural regeneration have been identified^3,4^, a study describing the stepwise morphogenetic changes that take place throughout a complete CNS regeneration is lacking. Utilizing the highly regenerative tunicate model *Polycarpa mytiligera*^5^, we characterized the morphological, cell proliferation, and transcriptomic dynamics that lead to entire CNS regeneration. The regenerated CNS of adult *P. mytiligera* expressed key neurodevelopmental markers that are not otherwise present in the adult CNS. Removal of the entire CNS resulted in high cell proliferation in the regenerated area. Transcriptome analysis revealed enhanced stem-cell related gene activity, with high expression of P53 and piRNA pathways preceding the activation of Notch, Wnt, and Nanos pathways. The CNS regeneration atlas created here depicts the transcriptomic landscape of the entire CNS regeneration process, revealing the core pathways that regulate neuronal response to injury, and the regeneration stage at which they are most pronounced. The molecular and cellular mechanisms controlling regenerative capacity that this atlas reveals could be used to develop approaches to enhancing neurogenesis in closely-related chordate species, including humans.

## Main

Adult mammals are unable to replace or repair lost neurons and damaged axons in their central nervous system (CNS)^2,3^. Consequently, CNS injuries and neurodegenerative disorders drastically reduce life quality and often lead to severe or even fatal outcomes. Our understanding of the mechanisms that enable neural regeneration is derived from invertebrates and non-mammalian vertebrates that are capable of recovering from severe CNS injury^3,6^. Tunicates are invertebrate chordates, a sister group of vertebrates, that share structures and cell types considered to be homologous to those in vertebrates’^7–9^, and are used as model systems for chordate development and regenerative studies^10–12^. In addition to providing insights into the evolutionary origin of the chordate CNS, the tunicate ability to regenerate their CNS may lead to the discovery of key cellular and molecular components of regeneration that are also likely to be found in vertebrates^11,13,14^. Here, we established the solitary tunicate *Polycarpa mytiligera* as a novel model system for regeneration and developmental studies, and characterized its extensive and unique regeneration abilities. Leveraging *P. mytiligera’s* extreme regeneration capacities^5,15,16^, we combined morphological and functional analyses with transcriptome sequencing in order to construct an integrated atlas of CNS regeneration. Comparing juvenile and adult CNS, we found that age has a significant effect on regeneration capacity, with juveniles demonstrating accelerated nerve regeneration abilities. Using an *in-vivo* EdU assay, we tracked cell proliferation patterns throughout the regeneration process and demonstrated the contribution of unspecialized dividing cells to the regenerating brain. We divided the regeneration process into three stages and characterized the differentially-expressed genes associated with each stage. We revealed that the re-establishment of the nervous system, by means of the newly-formed neural progeny, was associated with the expression of 239 conserved pathways, reflecting enhanced stem-cell related gene activity.

### Age-related variation in CNS anatomy and regeneration abilities

The adult ascidian CNS, composed of a single ganglion (brain^17^) and connected by a network of nerves to the different body parts, has an important role in controlling movement and behavior^18^. To achieve an accurate description of the morphological and cellular events underlying *P. mytiligera* CNS regeneration, a detailed description of the CNS anatomy of juvenile and adult animals was acquired using naive animals (Fig. 1). Serial sections and immunohistochemical staining revealed a similar morphology in both life stages. *P. mytiligera*’s brain consists of an outer cortex and a central neuropil (Fig. 1d, h). Nerve cell bodies are present in the cortex, surrounding a medulla where synaptic connections can be seen (Fig.1h). Four main anterior nerves, two posterior, and one ventral nerve sprout from the brain (Fig. 1f-g). The anterior nerves innervate the incurrent siphon and the anterior body wall, while the posterior nerves innervate the excurrent siphon and the remaining body wall. The ventral visceral nerve extends over the branchial basket toward the digestive tract (Fig. 1g). The brain is located adjacent to the neural gland (Fig. 1), a non-nervous structure that shares with the brain a common embryonic origin^19^. Together, the brain and the dorsal neural gland comprise the neural complex. The number of neural gland apertures opening into the pharynx varies between juveniles and adults. Whereas in juveniles, a single ciliated opening is observed (Figs. 1e-g), in the larger adults the duct splits into numerous ciliated openings (Fig. 1a, c). Both young and old animals were able to survive and regenerate following CNS removal. However, the time needed to complete a successful regeneration strikingly differed between the two age groups. Young animals regenerated their entire CNS within 7 days, while older animals needed 3 weeks to complete the same process (Fig. 2a). Interestingly, the newly-formed neural gland of the adult resembled in structure that of the juvenile. While the control adult gland has numerous openings (Figs. 1a, c), at 21 days post amputation (dpa), the regenerated neural gland had only a single detectable aperture (Fig. 2b), resembling the juvenile structure. This finding indicates the reactivation of embryonic developmental programs during adult regeneration^20,21^. Our morphological description of *P. mytiligera* CNS presents a simple brain composed of tubulin positive nerve cells that interact via synaptic connections, representing a basic model comparable with other chordates (Fig. 1). Based on these results, we were able to monitor the CNS regeneration process, distinguish between distinct developmental stages, and determine its endpoint (Fig. 2a).

**Fig. 1:**
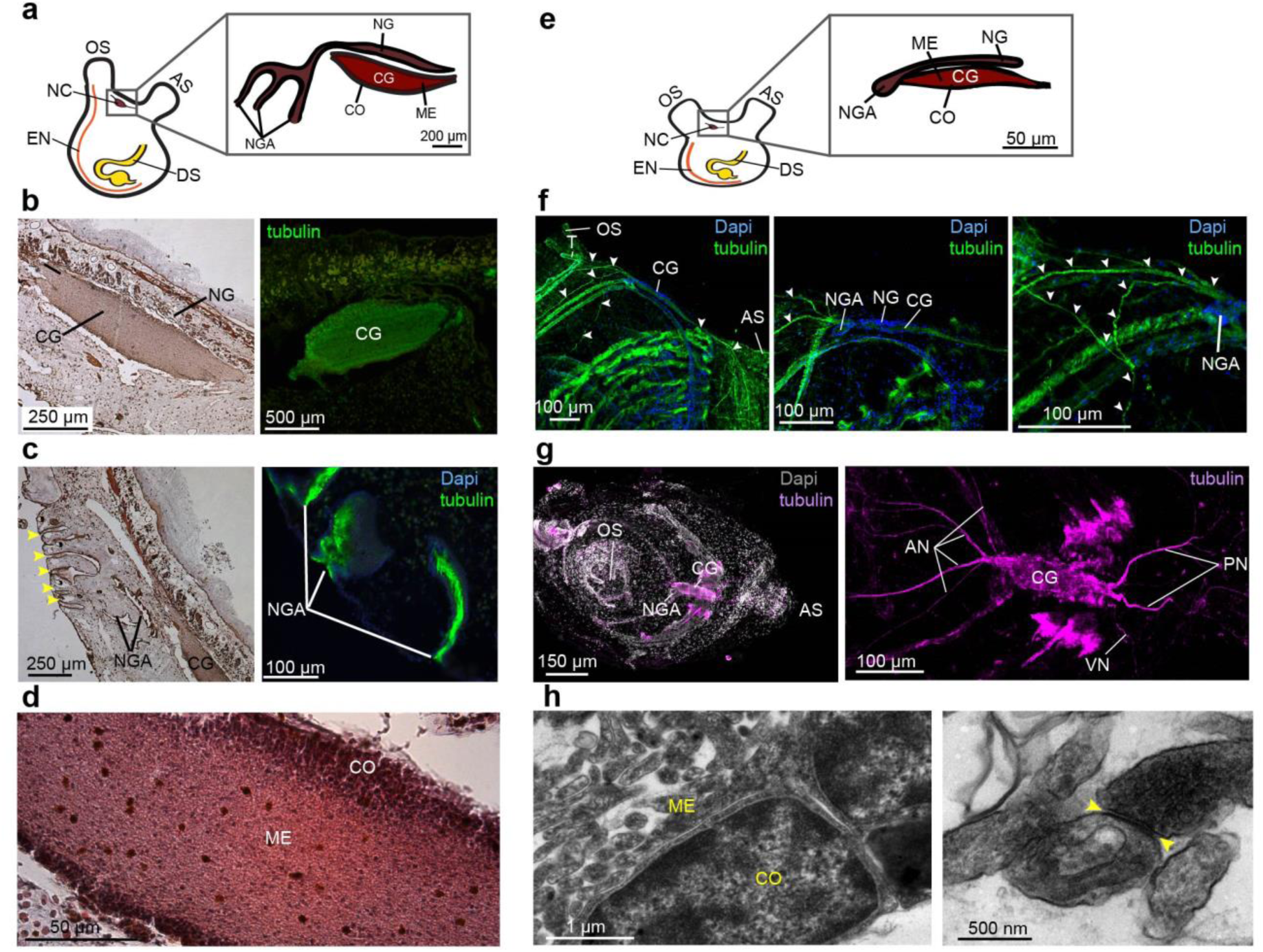
Comparative CNS morphology reveals minimal plasticity at different life stages. **a-d** , *P. mytiligera* CNS morphology during the adult life stage. **a**, Illustration depicting adult body plan, highlighting the CNS, located between the two siphons, and its components, the cerebral ganglion (CG), medulla (ME), cortex (CO), neural gland (NG), and neural gland aperture (NGA). **b**, Representative histological and immunohistochemistry staining of the cerebral ganglion. **c**, Representative histological and immunohistochemistry staining of the neural gland aperture. Neural gland ciliated apertures indicated by yellow arrowheads. **d**, Representative histological section of the cerebral ganglion. **e-h**, *P. mytiligera* CNS morphology during the juvenile life stage. **e**, Illustration depicting juvenile body plan highlighting the CNS, located between the two siphons, and its components, the cerebral ganglion and the neural gland with its single aperture. **f**, Whole mount immunofluorescence of *P. mytiligera* neural complex. Maximum intensity projections of confocal stacks. White arrows indicate nerve fibers. **g**, Whole mount fluorescent *in-situ* hybridization of tubulin showing the cerebral ganglion and associate nerves; anterior nerves (AN), posterior nerve (PN), and visceral nerve (VN). **h**, Transmission electron microscopy images of *P. mytiligera* neural complex showing the brain structures; medulla (ME), cortex (CO), and neural gland aperture (NGA). Synapse in the medulla is marked with yellow arrows. Atrial siphon (AS), digestive system (DS) endostyle (EN) and oral siphon (OS).

**Fig. 2.**
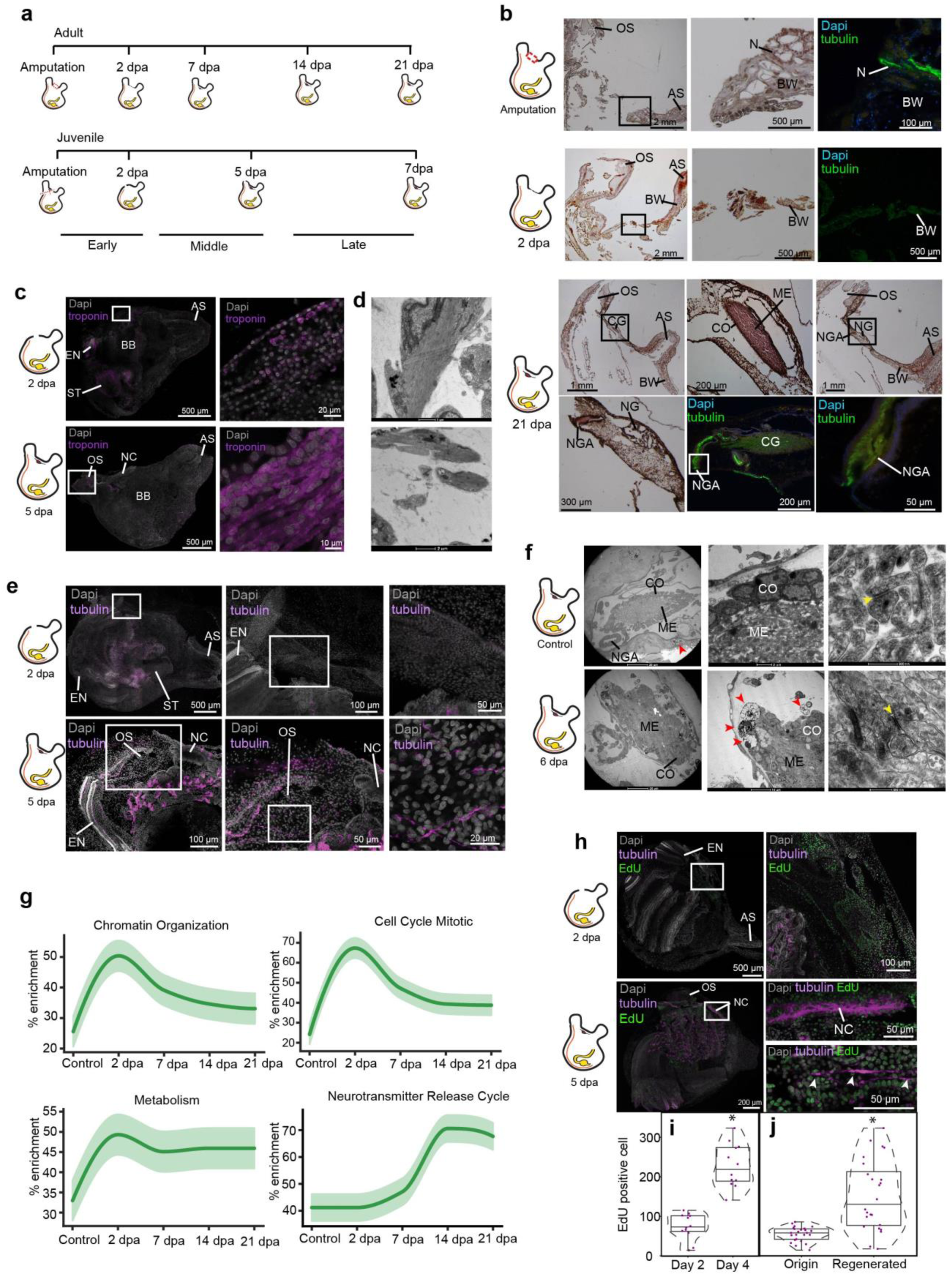
Three distinct anatomical and functional stages characterize CNS regeneration. **a,** Regeneration timeline showing the difference in regeneration rate between adults and juveniles along the three stages of CNS regeneration. **b**, Adult animal histological section and immunofluorescent staining (tubulin) showing CNS regeneration process along different time points. Enlargements of the square area appear in the following image in the panel. The CNS is absent following amputation. By 2 dpa we see beginning of closing of the open wound followed by complete regeneration of the CNS by 21 dpa. Anterior siphon (AS) body wall (BW) cerebral ganglion (CG), cortex (CO), medulla (ME), nerve fiber (N) neural gland (NG), neural gland aperture (NGA), and oral siphon (OS). **c,** Whole mount of regenerating animals showing troponin expression in regenerating muscle fibers at 2 and 4 dpa. **d**, Transmission electron microscopy images of regenerating muscle fibers at 6 dpa. **e**, Whole mount of regenerating animals showing tubulin expression in regenerating nerves fibers at 2 and 4 dpa. **f**, Transmission electron microscopy images showing the brain structures and cellular components in control (non-amputated) and 6 dpa animals. Synapse in the medulla is marked with yellow arrows. Immune cells are marked with red arrows. **g**, Gene enrichment plot of regeneration-associated and CNS related gene-sets during CNS regeneration. Light shaded regions indicate the 50% and 99% confidence intervals under a hypergeometric model. **h**, Whole mount staining of CNS regeneration at 2 and 4 dpa showing newly divided EdU positive cells in the regenerated tissue and in regenerating nerves fibers expressing tubulin. **i-j**, Quantification of EdU-positive cells in 100µm^2^ sections (n=4 sections per animal). Violin plots display the number of EdU-positive cells in each section. Data are mean ± SE. *P* values determined by Mann Whitney Wilcoxon test and are indicated above each boxplot. **i**, Number of EdU-positive cells in the regenerating tissue along the different time points (*n* = 3 per time point). **j**, Number of EdU-positive cells in the origin and regenerated tissue at 2 and 4 dpa (*n* = 3 per time point).

### Unspecialized proliferating cells contribute to the regeneration of the CNS

To study the cellular and transcriptomic signatures associated with CNS regeneration, we compared CNS samples taken from control animals (to establish a basal state) with CNS collected from diverse time points along the regeneration process. These samples were processed using multiple imaging and labeling methods that allowed us to focus on key regeneration stages (Fig. 2). Corresponding adult CNS samples were taken for RNA sequencing and used to construct a morphological and transcriptomic atlas of the entire regeneration process. Based on these results, CNS regeneration was divided into three milestones—early-, middle-, and late-stage regeneration (Fig. 2a). The early stage constitutes a period when progenitor recruitment and activation take place but prior to differentiation into the regenerated CNS. The middle stage is when proliferation and differentiation of progenitor cells to nerve cells take place while the CNS is undergoing regeneration which continue during the late stage where the CNS regain the morphological structures and cellular distribution pattern of the control structure.

To better characterize the cells that had contributed to the regenerated tissues we used fluorescent in situ hybridization of tissue-specific markers and conducted EdU pulse and chase experiments (Fig. 2 and Extended data Fig. 3). Our results indicate a possible contribution of undifferentiated progenitor cells that enter the S-phase during the early regeneration stages, to the formation of the regenerated CNS. Following amputation of the CNS, only the isolated tubulin-positive nerve fibers in the surrounding tissue remained (Fig. 2b). At the early stage epidermal tissue started to regenerate and close the open wound (Fig. 2b). This tissue was composed of regenerating troponin-positive muscle fibers (Fig. 2c, d), and nerve fibers had not yet re-innervated the tissue (Fig. 2e). During the first 16 h post amputation (hpa) no significant difference in the number of EdU positive cells was found between non-regenerated and regenerated areas. At 48 hpa a higher number of proliferating cells were located specifically in the regenerated area, indicating a local increase in proliferation rate. At this time point, these dividing cells did not express CNS specific markers (Fig. 2h and Extended data Fig. 3).

During the mid-regeneration stage, the wound became fully healed, and the CNS regenerated, with newly-formed nerve fibers sprouted from the brain and continued along the regenerating tissue to the siphons (Fig. 2e). This stage was also characterized by the highest number of EdU-positive cells. Some of these cells were found embedded in the regenerated brain and nerve fibers and were tubulin-positive, indicating their neuronal identity (Figs. 2h-j and Extended data Fig. 3). These results indicate a recent differentoiation event, as these neural progenies acquire their identity as nerve cells at this time point, following proliferation during the early regeneration stage.

At the late regeneration stage, the *de-novo* brain morphological structures and cellular distribution pattern resembled the control structure (Fig. 2b). TEM analyses verified that the regenerated brain contained a cortex and a medulla with visible synapses. However, the brain morphology was still not identical to the control brain at this point, as the cortex layer was not yet well organized into distinct layers (Fig. 2f). In addition, we observed several immune cells embedded in the cortex layer (Fig. 2f), implying a possible contribution of the immune system to the regeneration process^22^.

### *De novo* transcriptome assembly and gene expression analysis

To characterized the molecular signature of neural regeneration we performed the first CNS regeneration-centered solitary chordate transcriptome assembly. Using differential expression analysis, we first identified CNS specific gene expression patterns (Supplementary Table 1), and then experimentally confirmed them, demonstrating the validity of our experimental design. The detailed assembly pipeline is presented in the Extended data Fig. 1a.

The generated dataset was then analyzed using over-representation (ORA) and functional enrichment (GSEA), to identify the key pathways that had been enriched during CNS regeneration in comparison to the control CNS (non-amputated brain) (Extended data Fig. 1a). Moreover, by combining the DEseq2^23^ and bioinformatic tools that we had developed (all vs. all analysis)^10^, we were able to build a detailed gene expression atlas of the differentially expressed genes for each regeneration stage (Supplementary Table 2). Compared to previous methods, which only support comparisons between two experimental groups, our method allowed us to uncover differentially expressed genes in all possible combinations between contiguous and individual groups, resulting in a hierarchy of differently-regulated experimental groups for each gene. This method allowed us to characterize specific stepwise expression changes that would not have been possible using previous methods. Based on these analyses, a binary gene-time expression matrix (with 1 indicating dynamically “high” expression and 0 indicating “low” or zero expression) was created for every expressed gene recorded along each time point. The binary pattern chosen for each gene was the one most supported by its specific gene hierarchy. Using this binary gene matrix, we created enrichment plots for gene sets associated with regeneration (Figs. 2g, 3a and Extended data Fig. 4b).

**Fig. 3:**
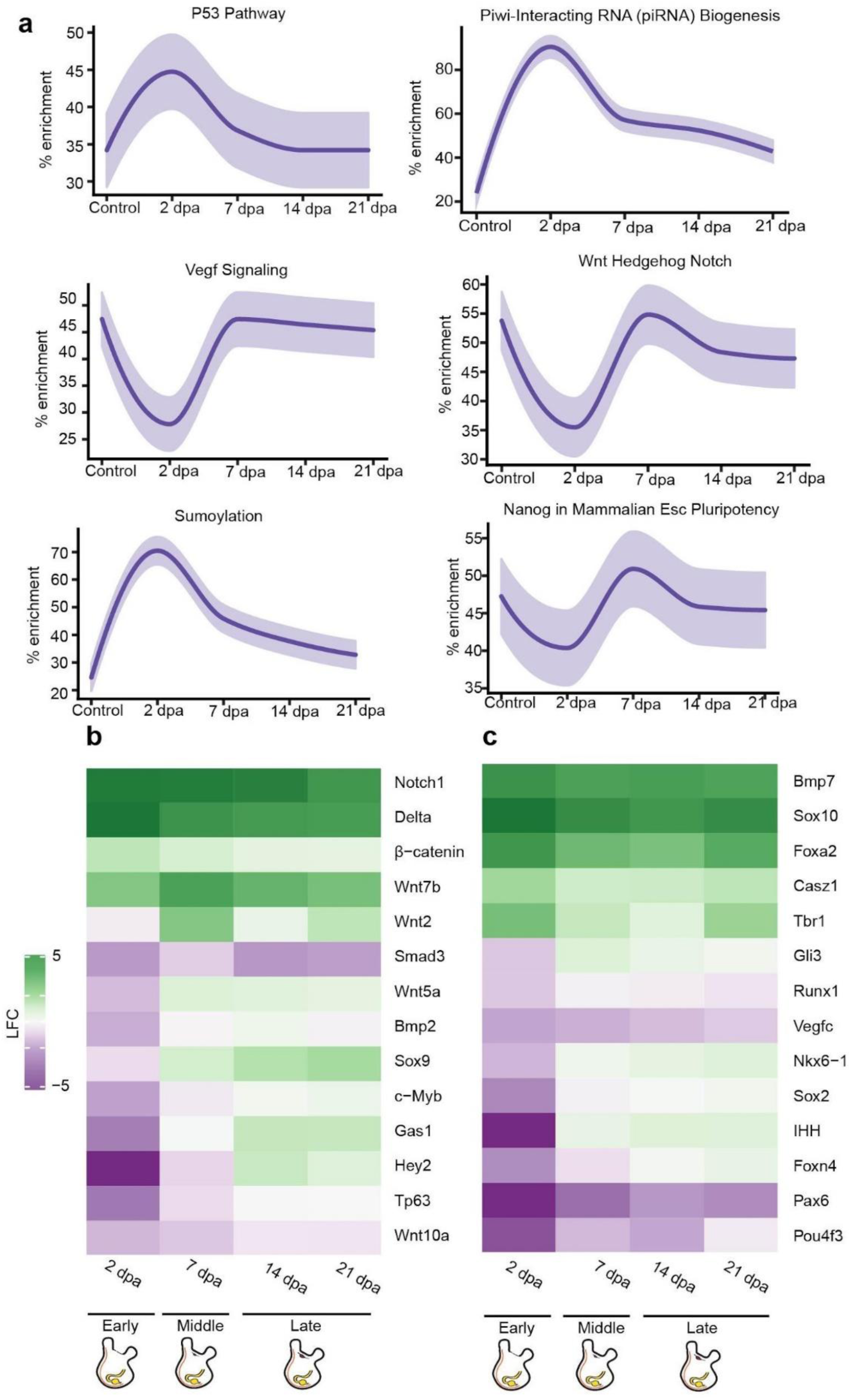
*De-novo* transcriptomic analysis of regenerating CNS reveals expression of stem cell markers and conserved regeneration associated transcription factors. **a**, Gene enrichment plot of regeneration-associated genes during CNS regeneration. Light-shaded regions indicate the 50% and 99% confidence intervals under a hypergeometric model. **b**, Heatmap of log fold change values for selected regeneration related genes significantly differ in comparison to the control. Green indicates a positive fold change (upregulated with respect to uncut CNS), and purple indicates a negative fold change (downregulated with respect to control). **c**, Heatmap of log fold change values for selected nervous system markers and regeneration related genes significantly differ in comparison to the control. Green indicates a positive fold change (upregulated with respect to uncut CNS), and purple indicates a negative fold change (downregulated with respect to control).

The genes specific to these time periods that possessed human or mouse gene homologs were analyzed using GeneAnalytics^24^, which allowed us to match gene lists with the relevant Gene Ontology (GO) and pathways (Supplementary Table 3).

### CNS regeneration is associated with enhanced stemness gene activity

To enable us to produce a detailed description of the transcriptomic landscape of CNS regeneration, we sought to determine the main regeneration-associated pathways that characterized each time point using our all vs. all analysis. The early regeneration stage was characterized by activation of cell proliferation and chromatin organization pathways (Fig. 2g), consistent with the results of our EdU assay. The main upregulated GO terms included cell signaling, nerve proliferation, and injury response pathways, reflecting the animal’s fast response to the amputation and damaged tissues (Extended data Fig. 2). Interestingly, this stage was enriched in several stem-cell related pathways (Fig. 3a), including the p53 pathway, a stress response pathway^25^ required for neurite outgrowth as well as for axonal proliferation and regeneration^26^. In the mouse brain, p53 is involved in the maintaining regenerative ability by regulating the proliferation of stem and progenitor cells^27^. It has been shown to exhibit several roles, including apoptosis, cell cycle arrest, and proliferation, depending on the different stages of the regeneration process^28^. In *P. mytiligera,* activation of this pathway occurs at the early regeneration stage and is inhibited at the later stages. These results indicate a negative regulation of neural stem-cell self-renewal and investment in specialization processes at the early time point, followed by stem cell maintenance as the CNS underwent reconstruction^29–31^. p53 transcriptional activities are primarily regulated through post-translational modifications, including sumoylation^32,33^. The sumoylation pathway was also enriched during the early stage of CNS regeneration (Fig. 3a). This pathway has a role in stem-cell proliferation^34^, and was shown to increase in mice following nerve lesion^35^. Another post-transcriptional regulatory pathway that was highly enriched following *P. mytiligera*’s CNS ablation was the Piwi-piRNA pathway (Fig. 3a). This pathway is involved in stem-cell maintenance in diverse organisms^36^, and was suggested to have an inhibitory role in neuron or Schwann cell responses during peripheral nerve injury in both nematodes and rodents^37^. Our results reveal an enrichment of this pathway following nerve injury, suggesting that it may play an important regulatory element during CNS regeneration. Further studies should be performed in order to determine the exact role of this conserved pathway in mediating axon regeneration following injury in *P. mytiligera*.

As regeneration progressed, dynamic changes in the enriched pathways could be observed. Upregulation processes associated with differentiation and tissue formation were expressed, as well as the enrichment of extracellular matrix organization processes (Extended data Fig. 2). Re-vascularization is an essential process during regeneration, requiring the re-growth of vessels and supportive cell types. Enrichment of Vegf signaling indicates such a healing process, consistent with the morphological description (Fig. 3a). At this stage the pathways associated with stem-cell activity that were detected at 2 dpa were replaced by a different set of stem-cell associated pathways, including Notch, Hedgehog, and Wnt (Fig. 3a), indicating a possible shift from stem-cell proliferation and differentiation to stem-cell self-renewal and maintenance^38–40^.

The middle and late regeneration stages showed an enrichment of the signaling pathways associated with tumor necrosis receptor factor 1 (TNFR1), fibroblast growth factor receptor (FGFR), and nerve growth factor (NGF) (Extended data Fig. 4b). These pathways play key roles in controlling the normal inflammatory and wound response to muscle and neuronal damage that is required to achieve regeneration^22,41,42^.

The late stage *de-novo* regenerated brain revealed an upregulation of GO terms such as “extracellular matrix constituent secretion” and “anterior/posterior pattern specification” (Extended data Fig. 2b), implying a continuous process of tissue assembly and specialization. Additional support for such a shift comes from the upregulation of voltage-gated ion channels genes, including *kcnh8* and *cacnb2* (Extended data Fig. 4a), and the enrichment of gene sets associated with the Nanos pathway (Extended data Fig. 4b) and neurotransmitter release cycle (Fig. 2g), reflecting morphogenesis processes and the establishment of functional synapses^43^. Moreover, the upregulation of interleukin−4−mediated signaling pathway at this time point (Extended data Fig. 2b) suggests a neurogenesis process, as this pathway plays a role in reducing proliferation and increasing neuronal differentiation^44^.

### Conserved molecular mechanisms underlie *P. mytiligera* CNS regeneration

Next, we sought to identify key regulatory factors that were differentially expressed during regeneration in relation to the control brain, and determine exactly when their expression was enhanced or inhibited along the regeneration process (Figs. 3b, c, Extended data Figs. 4, 5). Our results revealed a dynamic expression of transcription factors related to neuronal differentiation and neural cell fate specification (Figs. 3b, c, Extended data Figs. 4, 5). Among this gene list *sox10*^45^*, tbr1*^46^ and *pax-6*^47^ were differentially expressed following CNS removal. Fluctuation in expression patterns of these markers has been previously shown to mediate CNS regeneration in other chordates. In rats, upregulation of *sox10* has a role in differentiation and maintenance of stem-cell properties in the neural crest^45^, while low expression of *pax6* in zebrafish and lizards is suggestive of axon regeneration^48^. These results indicate a possible conserved role of these factors during nerve regeneration in *P. mytiligera*, as both s*ox10* and *tbr1* were upregulated throughout the entire process, while *pax-6* was consistently down-regulated.

The nervous system development genes, *foxa2*^49^ and *six1*^50^, were differentially expressed during CNS regeneration in relation to the control CNS (Figs. 3b, c, Extended data Figs. 4, 5). In mice, *foxa2* is required for specification and differentiation steps of neurons at progressively higher doses in order to facilitate differentiation^51^; while *Six1* expression is required for the proliferation of neuroblast progenitors^50^. Our results show a similar trend of these factors, as both *foxa2* and *six1* were upregulated during *P. mytiligera* CNS regeneration.

Several neurogenic transcription factors were also differentially expressed during the regeneration process. Both *c-myb* and *sox2* were upregulated during the late stage of CNS regeneration. *Sox2* is a universal neural stem-cell marker used to identify cells with self-renewal ability and multipotent differentiation found in both embryonic and adult animals^52^. In the adult brain *sox2* is involved in the proliferation and maintenance of neural stem cells as well as in neurogenesis^53^. The hematopoietic transcription factor and proto-oncogene *c-myb* regulates neural progenitor proliferation and has an important regulatory role in the neurogenic niche of the adult mouse brain. A previous study has shown that *c-myb* expression declines when neural stem cells differentiate^54^. Indeed, *c-myb* is suggested to regulate both *sox-2* and *pax-6*gene expression to maintain cell cycle progression and neuroblast generation^54^. The expression of these neural stemness markers following CNS removal, strongly suggests the involvement of stem cells during regeneration in *P. mytiligera*.

As noted in the previous section, our data revealed the dynamic expression of highly conserved signaling pathways, such as Wnt, Notch, and Hedgehog, all specifically enriched at the middle and late stages of regeneration (Fig. 3a). Closer observation of the gene sets composing these pathways revealed potential key regulators of CNS regeneration (Figs. 3b, c, Extended data Figs. 4, 5). The Wnt/β-catenin pathway is represented in *P. mytiligera* by different factors, including *wnt7b, wnt5a,* β-catenin, low-density lipoprotein receptor-related protein 5 (*lrp5*), and *lrp6*, all of which show a dynamic expression pattern along the regeneration process (Figs. 3b, c, Extended data Figs. 4, 5). Components of the Notch signaling pathway were also differentially expressed during *P. mytiligera* CNS regeneration. Our results revealed upregulation of the receptor protein gene *notch-1* and its ligand, *delta,* down-regulation of the signal mediator *rbpj*, and dynamic expression of a downstream transcriptional target *hey-2* (Figs. 3b, c, Extended data Figs. 4, 5).

Our transcriptome results emphasize the involvement of these conserved pathways at the late stage of CNS regeneration. As both pathways have vital functions during embryonic development, tissue regeneration, and stem-cell maintenance in a variety of model species, including vertebrates^55–59^, their dynamic expression in *P. mytiligera* provides an opportunity to further study their involvement in neuron regeneration in a simple and accessible new chordate model system.

## Discussion

Here we combined morphological analysis with transcriptome sequencing in order to construct an integrated atlas of CNS regeneration in an emerging chordate model system. The transcriptomic and cellular signatures associated with CNS regeneration evolves over time points that represent early, mid, and late regeneration stages (Fig. 4). In the early stage, extensive proliferation of progenitor cells occurs at the regenerating area while many genes reflect a wound response by facilitating cell division and stress signals (including *gas-1*, *cdc-42* and *hsp90*). At this time point, two key pathways associated with stem cells; Piwi interacting RNA biogenesis and P53, are activated (Fig 4). As regeneration progresses these pathways are inhibited and conserved pathways associated with regeneration and stem-cell self-renewal are expressed (including Wnt, Hedgehog, and Notch), as well as candidate homologous genes associated with neuronal tissue growth and differentiation (for example, *sox-2* and *pax-6*) (Fig. 3). These results aligned with the cellular data as we see newly-divided cells that acquire their neuronal identity at this time-point. The late stage of regeneration is characterized by the activation of genes related to functioning synapses and the maintenance of neuronal stem cells. Thus, within a three-week time frame, classical neurodevelopmental events occur that facilitate CNS regeneration (Fig 4).

**Fig. 4:**
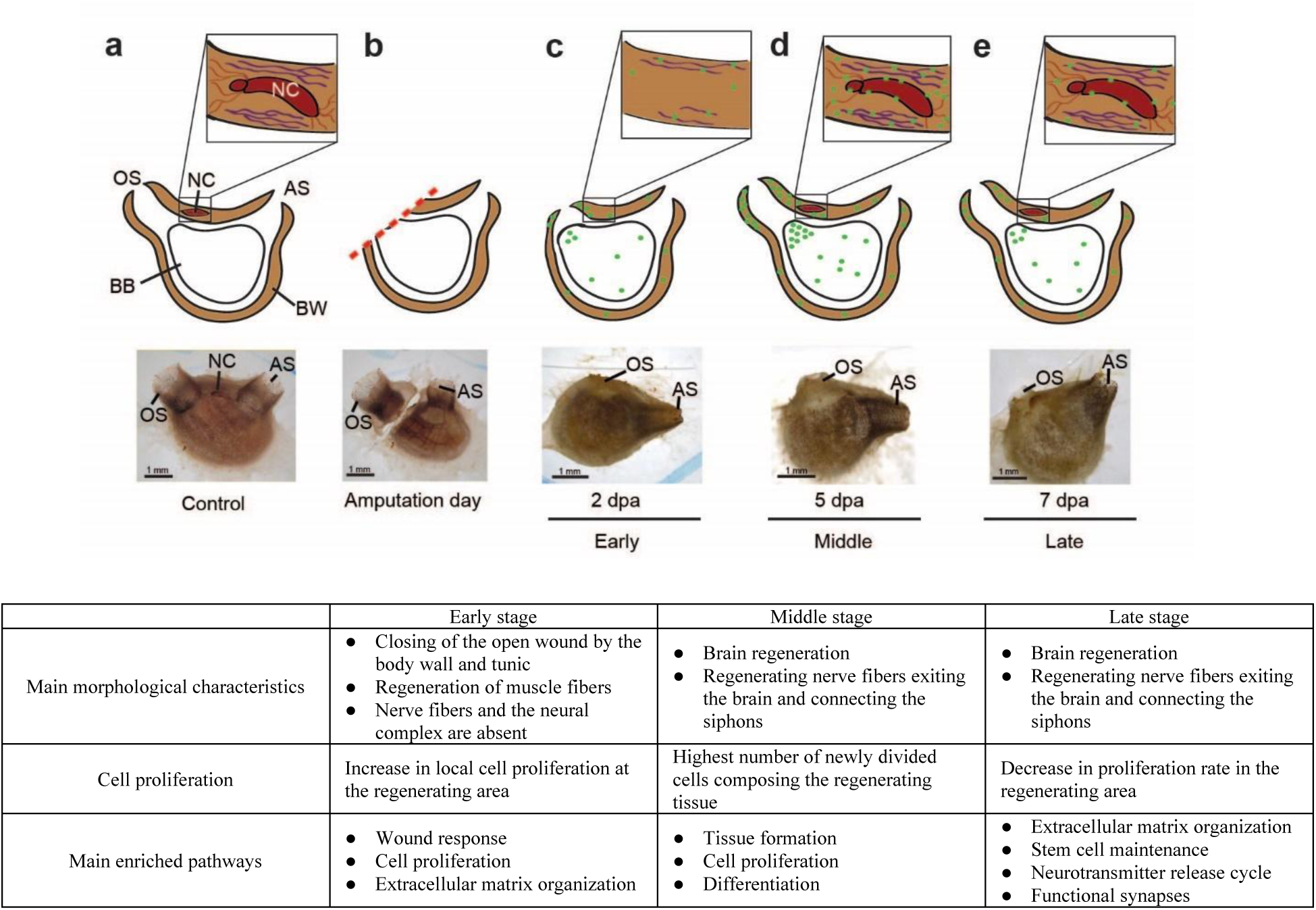
The cellular and transcriptome blueprint of CNS regeneration. **a-e** , Summary of the main cellular events leading to CNS regeneration (green-EdU positive cell, red-nerve fibers, and purple-muscle fibers). **a**, Illustration and *in-vivo* image showing a control (pre-amputation) animal. The brain (neural complex, NC) is embedded in the body wall (BW). Nerve fibers connecting the brain with the surrounding tissue. Muscle fibers embedded in the body wall, surrounding the brain. Atrial siphon (AS), branchial basket (BB), and oral siphon (OS). **b**, Illustration and *in-vivo* image showing an animal immediately following amputation. **c**, Illustration and *in-vivo* image showing an animal at early stage of regeneration (2 dpa). The open wound is closed by epidermal tissue composed of newly regenerating muscle fibers. The brain and associated nerves are absent. Higher levels of EdU positive cell embedded in the regenerating tissue in comparison to non-regenerating tissue. **d**, Illustration and *in-vivo* image showing an animal at middle regeneration stage (5 dpa). The open wound is closed by epidermal tissue composed of regenerating muscle fibers. Nerves fibers appear at this stage and are connected to the regenerated brain. At this point we see the highest level of EdU positive cells embedded in the regenerating tissue. **e**, Illustration and *in-vivo* image showing animal at late regeneration stage (7 dpa). At this point we see a decrease in the level of EdU positive cells embedded in the regenerating tissue, albeit still higher than the level found in non-regenerating tissue. **f**, Summary table of the main morphological, functional, and molecular processes involved in *P. mytiligera* CNS regeneration.

Previous studies have demonstrated the crucial role of stem-cells in colonial ascidian regeneration and asexual development processes^60–62^. Since brain regeneration initiate the proliferation of progenitor cells and triggers the expression of known stem-cell transcriptional regulators, including *sox2*, *hes1*, and *c-myb* (Figs. 3), we suggest that CNS regeneration in *P. mytiligera* is mediated by a population of adult stem cells. As the efficiency of axon regeneration can be experimentally manipulated^3^, the identification of regeneration-associated genes offers a promising approach for developing therapeutic applications *in vivo*.

Our study presents a detailed road map of complete CNS regeneration in a solitary chordate, in which the activation and inhibition of candidate neurogenesis factors are documented (Supplementary Table 3). Analysis of these transcription factors in this highly regenerative emerging model system, and comparisons of their expression pattern with other closely-related species with varied regenerative capabilities, will enable a better understanding of the evolution of the regeneration mechanism and the identification of targets that can enhance regenerative capacity in other chordates, including in humans.

## Methods

### Experimental Design

Our main research aim was to provide a detailed description of CNS regeneration in a new chordate model system. To this end we characterized the morphological, functional, and molecular processes underlying *P. mytiligera* CNS regeneration.

#### Morphological analysis

We first described the control CNS in juvenile and adult animals to better understand this system anatomy and to determine the end point of the regeneration process. Next, we monitored the reconstruction of the CNS using various methods, including histology, immunolabeling, and *in-situ* hybridization at different time points following CNS removal (Fig. 1). Our findings enabled us to divide the regeneration process into three key stages that reflect similar morphological criteria in both age groups (Fig. 2a).

#### Functional analysis

Cell proliferation underlies the formation of new tissues in diverse regenerative organisms, including *P. mytiligera.* Our goals were therefor to better characterize the identity of post-amputation dividing cells and to study their involvement in nerve regeneration. First, we sought to describe the location of the proliferation events and determine whether CNS removal resulted in local or systemic cell proliferation. Next, we sought to monitor the identity of the proliferating cells using the tissue-specific markers alpha-tubulin (nerves and cilia^63–65)^ and troponin (muscle^66^). To achieve these goals, we applied the EdU proliferation assays to regenerating juveniles and monitored the location of cells that entered the S-phase following CNS removal. We then analyzed the expression of tubulin and troponin in these dividing cells to determine the regenerative stage at which they acquire their identity (Fig. 2h, Extended data Fig. 3).

#### Molecular analysis

We performed *de-novo* transcriptome analysis using CNS samples of adult animals from the three chosen regeneration stages, in order to analyze the molecular landscape of neural regeneration. As adult stem cells have been shown to mediate whole body regeneration in other tunicates, we hypothesized that CNS regeneration would involve the activation, proliferation, and specialization of a similar population in our highly regenerative model. Focusing on regeneration and stem cell markers, we compared the expression patterns of each regeneration stage with the control and with all other time points, in order to identify the enriched pathways and key transcription factors activated or inhibited during CNS regeneration (Figs. 2g, 3 and Extended data Figs. 4, 5). This comprehensive analysis provided a valuable insight into the factors and pathways that conduce to wound healing and neurogenesis, as revealed by our new chordate model system.

### Animal collection and culturing

During March-April 2018 *P. mytiligera* adult individuals were collected from the Gulf of Aqaba (Eilat), Red Sea (Israel). The animals were maintained at the Eilat Inter-University Institute (IUI) in aquaria with running seawater for four days of acclimation prior to the onset of the experiments. *P. mytiligera* juveniles were obtained from a breeding culture established in the IUI as previously described^67^.

### Regeneration experiment

For the morphological description of the regeneration process, and selection of representative time points, adults (n=24) of similar size range (4±1 cm, from the tip of the oral siphon to the base of the animal) were anesthetized^68^ and dissected. The CNS was removed using a scalpel. Both treated and non-amputated (control) animals (n=4) were maintained in running seawater for the duration of the experiment. Response to touch was used to determine survival. Juveniles of the same age (3 months old, n=24) were dissected and their CNS was removed. Due to their small size the oral siphon and the CNS were amputated together to ensure removal of the entire CNS.

### Histology and immunohistochemistry

Regenerating adult animals (n=20) were fixed at different time points along the regeneration process (amputation day, 2-, 7-, 14-, and 21 post-amputation). Following relaxation with menthol, the animals were fixed in 4% formaldehyde in seawater partly separated from the tunic, and dehydrated with ethanol at increasing concentrations prior to paraffin (Paraplast Plus, Lecia) embedding. Sections (7μm) were mounted on glass slides, deparaffinized, and stained with standard hematoxylin and eosin solutions.

For immunolabeling, chosen sections were rehydrated in xylene and graded series of ethanol to phosphate-buffered saline (PBS). Juveniles (n = 10) were fixed in 4% paraformaldehyde in phosphate-buffered saline (PBS) overnight at 4°C. Fixed juveniles were then washed three times (5 min each) in PBS followed by a 10 min wash in TritonX-100 at room temperature (RT). Samples were blocked in PBT + 3% BSA for 3 h at RT. Antibodies were added directly to the blocking solution overnight at 4°C. To visualize nerves and cilia, we used a mouse monoclonal anti-acetylated tubulin antibody (Sigma T7451) diluted 1:1000 in blocking solution^63,65,69^. Samples were then washed twice for 10 min each in PBT. Secondary antibodies (ThermoFisher, Alexa Fluor goat anti-mouse IgG 488 A-11001) were added at 1:500 dilution to the blocking solution for 3 h at RT. Samples were then washed in PBS three times for 10 min each. Samples were stained with DAPI (Sigma, 1 μg/ml in PBSTx) and mounted. Images were taken using a fluorescent microscope.

### Transmission electron microscopy (TEM)

For a detailed description of *P. mytilgera* CNS morphology, juveniles (3 months old, n=5) were fixed in 1.5% glutaraldehyde buffered with 0.2M sodium cacodylate, pH 7.4, plus 1.6% NaCl.

After washing in buffer and post-fixation in 1% OsO4 in 0.2M cacodylate buffer, the specimens were dehydrated and embedded in epoxy resin (Sigma-Aldrich). Ultra-thin sections (80 nm thick) were stained with uranyl acetate and lead citrate to provide contrast. Photomicrographs were taken with a FEI Tecnai G12 electron microscope operating at 100 kV. Images were captured with a Veleta (Olympus Soft Imaging System) digital camera.

### An RNA-Seq Catalog for central nervous system Regeneration

To characterize the molecular events that take place during *P. mytiligera* regeneration, and to establish a reference map of CNS regeneration amenable to inter-species comparison, we profiled 12 samples of CNS tissue and surveyed gene expression changes. CNS samples (n=3) from untreated animals were used to reflect the start and end points of regeneration. Based on these morphological features, pools of regenerating CNS tissues were collected at five time-points: 2 days (n=2), 7 days (n=2), 14 days (n=3), and 21 days (n=2) post-amputation, to provide an informative perspective on the molecular events leading to CNS regeneration.

### RNA-Seq, read mapping, and transcriptome assembly

Samples were prepared at the IUI during September 2017. Fifteen *P. mytiligera* individuals (6±1 cm, from the tip of the oral siphon to the base of the animal) were used for neural complex regeneration experiments. Neural complex tissue samples were prepared for RNA extraction (RNAeasy mini kit from Quiagen) at different time points along the regeneration process. Total RNA was extracted following the manufacturer’s protocol. cDNA libraries were then prepared from high-quality samples (RNA integrity number (RIN) > 8) at the Weizmann Institute of Science, Life Sciences Core Facilities, Israel. Barcoded library samples were sequenced on an Illumina NextSeq 500 (2 × 150 bp) at Stanford University, CA, USA.

*De-novo* transcriptome assembly using the Trinity pipeline (https://informatics.fas.harvard.edu/best-practices-for-de-novo-transcriptome-assembly-with-trinity.html) was followed (Supp. 1): Adapter sequences and low-quality bases were trimmed using fastp^70^. Sequencing errors were corrected using Rcorrector^71^ (v1.0.4) and all reads that were not correctable were discarded. Reads were aligned to the known contaminant *phiX* (Illumina igenome) and known rRNA sequences^72^ using Bowtie2^73^ (2.4.1) with the setting-very-sensitive-local. Aligned reads were removed. Remaining reads were assembled into transcripts using Trinity^74^ (v2.9.1). Assembly quality for each transcript was assessed using TransRate^75^ (1.0.3). Only transcripts with p_good >0 score were used for further analysis. TransDecoder^74^ (v5.5.0) was used for identifying candidate coding regions within the transcript sequences. Homologous sequences were identified and annotated, based on the candidate coding region transcripts, using BLASTP (NCBI), cut-off 1e-10, to align with (1) the SwissProt protein database; and (2) solely on the *Mus musculus* curated SwissProt database (downloaded on 21-02-2020).

Salmon^76^ was used for quantifying gene expression with the following settings: --validateMappings, -- numBootstraps 100, --seqBias and –gcBias. Transcript-level abundance were imported using tximport^77^ (v1.16.1) to DESeq2^23^ (1.26) for gene-level analysis. Genes with ≤5 supporting reads in ≤2 samples were discarded. The DESeq2 analysis was employed using both the likelihood ratio and Wald tests, to identify all differentially expressed genes between the sequential regeneration stages (FDR<0.05).

Sets of genes were tested for enrichment of Gene Ontology (GO Biological Process, Molecular Function and Cellular Component) terms. For a set of genes with significant up and down effects, found using the Wald test, an over-representation analysis (ORA) was performed using the enricher function of clusterProfiler^78^ package (v3.16.1) in R with default parameters. A gene set enrichment analysis (GSEA) was also performed on the entire assembly using a scoring based on the log fold change of each gene on its respective time point using the GSEA function of clusterProfiler^78^ package (v3.16.1) in R with default parameters. In both cases Significant GO terms were identified with an FDR < 0.05.

The gene ontology terms were obtained from a manually created database based on the SwissProt curated *Mus musculus* GO annotations, using the makeOrgPackage function of AnnotationForge^79^. Pipeline summary is presented in Extended data Fig. 1.

### Identification of regeneration-associated differentially expressed genes

The generated dataset was analyzed using two methods:

1. Identification of differentially expressed genes between each time point and the control homeostatic CNS sample. For each time point, transcript levels were compared between each pool of regenerating fragments and the control sample (Supplementary Table 1).
2. Identification of differentially expressed genes between all possible combinations of all time points. The identification of chronologically differentially-expressed genes and the formation of binary tables have been described previously^10^. Briefly, we used DEseq2^23^ (*REF*; FDR < 0.05) to identify the differentially-expressed genes between all possible combinations of contiguous and individual time points, resulting in a hierarchy of up-and-down regulated time points for each gene for each age group. For each gene in each experimental group its specific time point hierarchy was then used to score all the possible binary patterns, with each pattern’s score being the number of shared up-and-down regulated time points between it and the hierarchy, while subtracting the number of up-and-down regulated time points of which the pattern and hierarchy disagreed. The pattern with the highest score was used for that gene and regeneration stage pairing. Based on these analyses, a binary gene-time expression matrix for every expressed gene recorded along the time points was produced, with 1 indicating dynamically “high” expression and 0 indicating dynamically “low” expression (Supplementary Table 2). Using the binary matrix, we identified pathways and Go terms associated with each time point (Figs. 2d, 3a and Extended data Fig. 4b, Supplementary Table 3).

### Gene enrichment plots

Gene enrichment plots have been previously described in^10^. Briefly, at each time point the proportion of genes in a gene set that are active (indicated by a 1 in the gene-time expression binary matrix defined above) is calculated. This gives a value between 0% (no genes in common) and 100% (all genes in the gene set are active at that time). A baseline expectation of the proportion of overlapping genes is calculated using a hypergeometric model that gives the likelihood that the same number of genes as in the selected gene set would be randomly selected from the matrix. In addition, the 68% confidence interval (1 standard deviation) of the proportion of shared genes (‘enrichment’) from the hypergeometric model is calculated and plotted, and presented as a shaded region in the plot. The baseline is then subtracted from the values calculated, with the confidence interval also subtracted, to present the expected range of values and the extent to which the actual enrichment result differs from a null result. If the baseline expectation is greater than the actual enrichment (i.e. the subtracted value will be negative) a value of 0% is used (as a negative percentage is considered meaningless).

### Use of specific pathway gene sets

For the gene enrichment plots the following gene sets were used: For the pathway and go terms identified by GeneAnalytics within our time data, we focused on the gene sets of the same names from PathCards. From each of these gene sets we removed any gene that did not have a putative homolog to a known *Polycarpa* gene. Consequently, the percentages within the enrichment plots refer to these curated *Polycarpa* specific gene sets. If a gene name appeared more than once in the *Polycarpa* gene model annotation, all matching *Polycarpa* gene ids were included in the gene set.

### Gene cloning and transformation

Gene-specific primers were designed from the transcriptome sequence and synthesized by Integrated DNA Technologies (IDT). Their oligonucleotide sequences were as follows: Troponin C 5’GGATTTGACGGAAGAGCAGA3’ and 3’TCATTGCCACGAACTCTTCA5’. Tubulin alpha-1A chain 5’CTGCAGACGAAACCTTGACA3’ and 3’TAAACCGTATCACCGTGCAA5’.

Genes were amplified from *P. mytiligera* cDNAs using gene-specific primers and cloned into pGEM-t vector using the manufacturer’s protocol (Promega; CAT #A1360). Vectors were transformed into *E. coli* Top10 by the heat-shock method. Briefly, 100 μl of bacteria were mixed with 5 μl of cloned vector, incubated on ice for 30 min, and then subjected to 42°C for 45 sec. The transformed bacteria were then supplemented with 350 μl of SOC medium, and following 1 h of recovery at 37°C, plated on agarose plates containing 1:2000 Ampicillin, 1:200 Isopropylthio-b-D-galactoside (IPTG), and 1:625 5-bromo-4-chloro-3-indolyl-β-D-galactopyranoside (X-gal). Colonies were grown overnight at 37°C, and screened by colony PCR using M13F and M13R primers with the following PCR program: a. 5 min at 95°C; b. 34 cycles of 45 sec at 95°C, 60 sec at 55°C, and 2 h 30 min at 72°C; c. 10 min at 72°C; d. hold at 10°C. Reactions were analyzed by gel-electrophoresis, and correctly-sized gene products were grown overnight in Luria Broth media (LB), supplemented with 1:2000 Ampicillin at 37°C. Plasmids were purified from overnight cultures using the NucleoSpin Plasmid Miniprep Kit (Macherey-Nagel; CAT #740588 cloned gene sequences were sequenced by Sanger sequencing.

### Fluorescent *in-situ* hybridization

Fluorescence *in-situ* hybridization (FISH) was based on the published protocol^80,81^ with some modifications (Supplementary item 1). Essentially, fixed animals were separated from their tunic and opened along the endostyle. Following bleaching and proteinase K treatments, samples were incubated with DIG/ DNP-labeled riboprobes overnight in hybridization solution, at 60°C. After hybridization, samples were washed twice in: pre-hyb solution, 1:1 pre-hyb-2X SSC, 2X SSC, 0.2X SSC, PBSTx. For the blocking step prior to antibody incubation, 0.5% Roche Western Blocking reagent and 5% inactivated horse serum in 1xPBSTx were used. Samples were then incubated with antibodies (anti-DIG-POD, Fab fragments) overnight at 4°C. Antibody solution was washed with 1xPBSTx. Rhodamine or FITC dyes were used in tyramide development. For peroxidase inactivation, samples were washed with 1% sodium azide solution for 90 min at room temperature. Finally, samples were counterstained with DAPI (Sigma, 1 μg/ml in PBSTx) for 1 h, mounted and photographed using a Zeiss LSM 880 scanning laser confocal microscope.

### *In-vivo* cell labeling experiments

Cell proliferation was detected by incorporating 5-ethynyl-2-deoxyuridine (EdU) into replicating DNA. *P. mytiligera* juveniles (3 months old) were divided into two groups, dissected, and separated from their neural complex and oral siphon using a scalpel (Extended data fig. 3a). One group (n=3) was exposed to a 16 h EdU pulse following amputation and was fixed immediately afterwards. The second group (n=9) was exposed to a 16 h EdU pulse 32 h following amputation and left in seawater to regenerate. Fixation was done at 2-, 5-, and 7-days post-amputation (n=3 per each time point) (Extended data Fig. 3a).

For the pulse experiments animals were incubated with 10 μmol/L EdU (Invitrogen, Carlsbad, CA) in 5 mL of MFSW for 16 h in Petri dishes. Following completion of the labeling, animals were fixed for 12 h in 4% FA, rinsed three times in 1 × phosphate-buffered saline (PBS), and processed for EdU detection using Alexa Fluor azide 488 at room temperature, according to the instructions of the Click-iT EdU Alexa Fluor High Throughput Imaging Assay Kit (Invitrogen). Samples were stained with DAPI (ThermoFisher 33342) (1 μg/mL in PBS) and mounted in VECTASHIELD (Vector Laboratories RK-93952-28) using coverslips.

### Statistical information

All results are expressed as mean ± SE. Statistical analyses were performed using R Studio. Statistical analysis was performed using two sided Mann Whitney Wilcoxon test throughout the study. Difference was considered significant as follows: *P < 0.05.

## Acknowledgments

We would like to thank Dr. A. Colorni and Ms. N. Paz for their editorial assistance. We are grateful to the Inter-University Institute (IUI) and the National Center for Mariculture (IOLR-NCM) staff for their ongoing support and use of the respective facilities.

We acknowledge the Yitzhak Navon Ph.D. scholarship to TG, The Erasmus Plus scholarships, the COST (European Cooperation in Science and Technology) Short Term Scientific Mission (COST Action MARISTEM – CA 16203), the Naomi Foundation and the SDB Emerging Models grant to TG, which allowed collaborative research between Tel Aviv University, the University of Padova and Stanford University.

## Author contributions

Conception and design: T.G., N.S., L.M., and A.V.; mariculture and sample collection: T.G.; RNA isolation: T.G; sequencing and transcriptome analysis: T.Z., Y.V., A.V., and M.K.; statistical analysis: Y.V., T.Z., A.V., T.G., and, M.K.; *in situ* hybridization preparation: T.G., N.H., and O.W.; light, electron, and confocal microscopy: T.G., L.M., O.W., and F.C.; writing of the manuscript: T.G., N.S., L.M., and A.V.

## Extended data

**Extended data Fig. 1.**
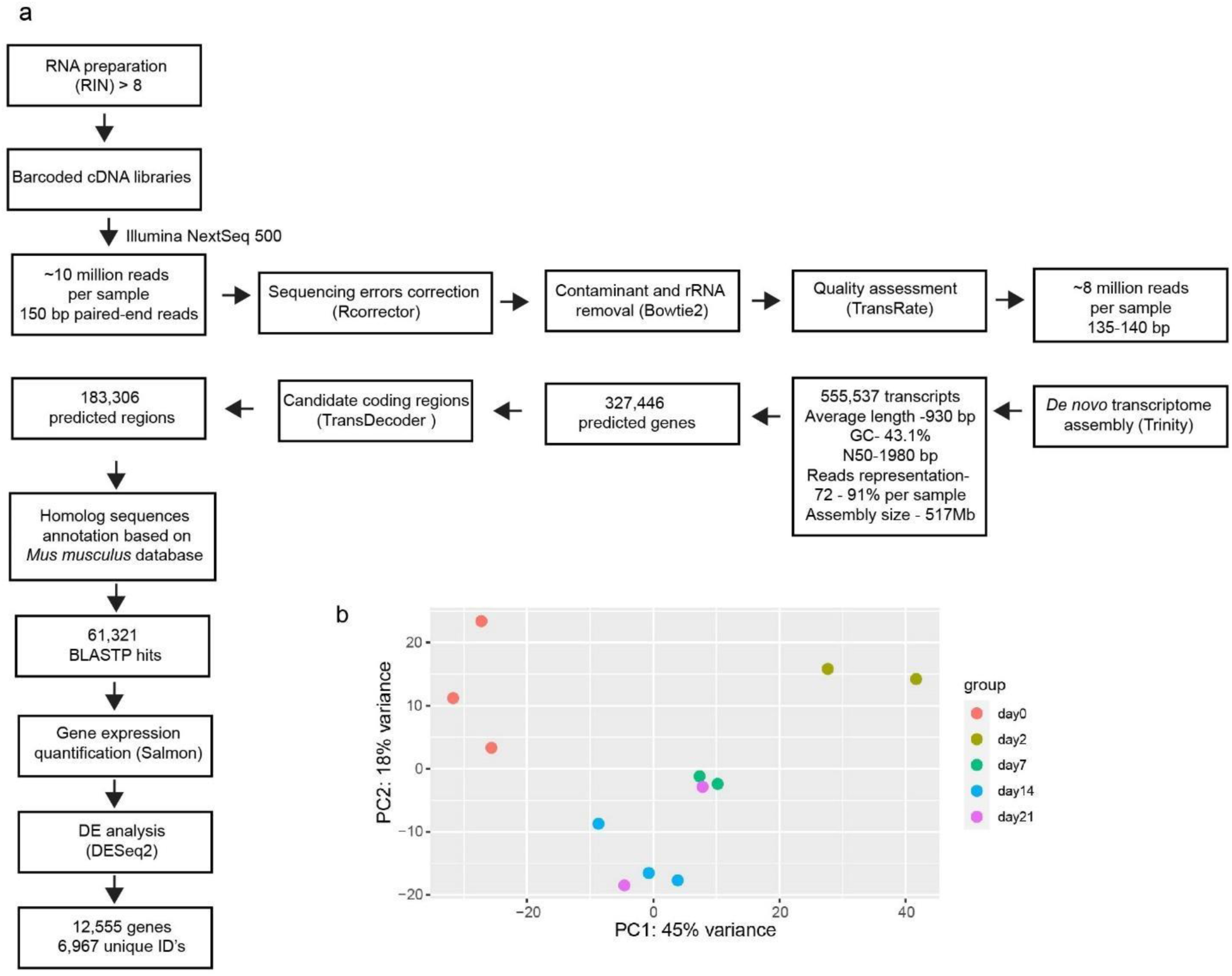
*P. mytiligera de-novo* transcriptome assembly summary. **a**, Pipeline for *P. mytiligera* RNAseq and *de-novo* assembly process. **b**, Principal component analysis (PCA). RNA-Seq data sets from control and different time points of CNS regeneration show separation between the CNS control sample and the regenerating samples, and between the early stage of regeneration (2 dpa) and late stages (7, 14 and 21 dpa), which are grouped together.

**Extended data Fig. 2.**
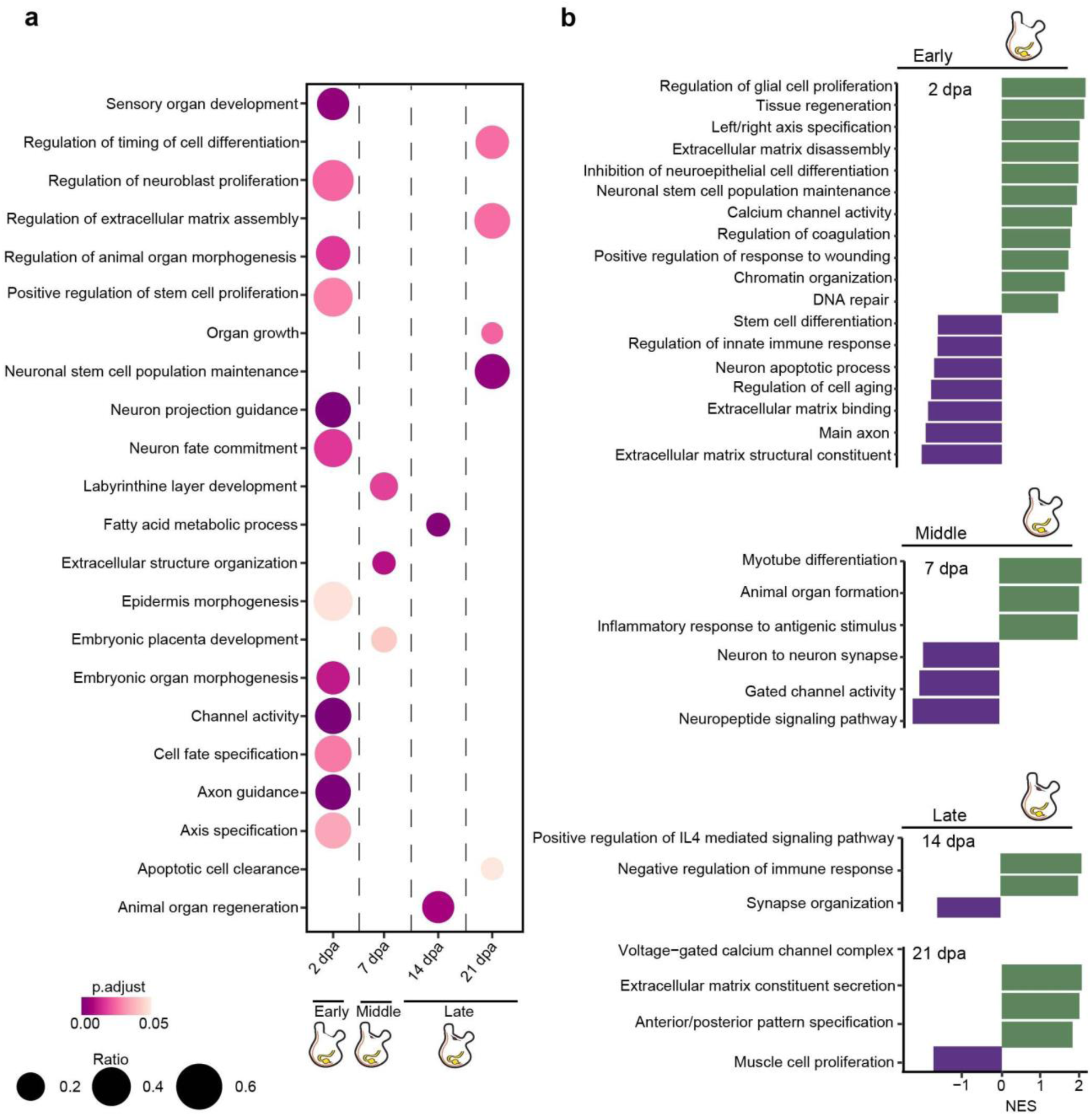
Gene ontology categories enriched at the different stages of CNS regeneration. **a,** Gene ontology (GO) term enrichments for each of the time points. The enrichment of each GO term is indicated by a circle, where the area corresponding to the fraction of genes annotated with that term is present in the cluster, and the color of the circle corresponds to the adjusted p-value of term enrichment (ORA, FDR≤0.05). **b,** Gene sets are plotted relative to normalized enrichment scores (NES). Categories with negative or positive NES are down- or upregulated, respectively (GSEA, FDR <0.5).

**Extended data Fig. 3.**
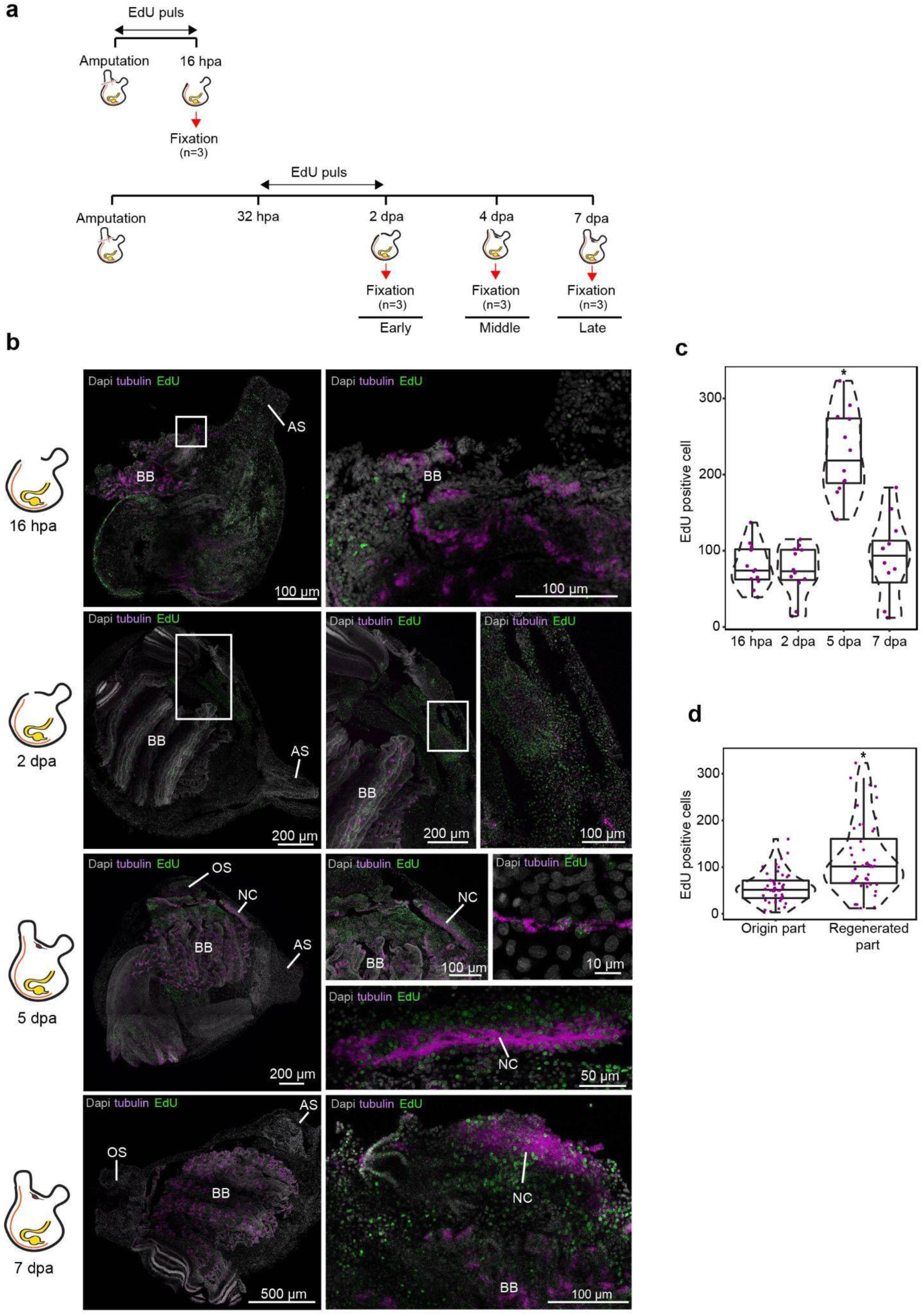
Cell proliferation increase during CNS regeneration. **a**, Experimental timeline and illustration of experimental paradigm. **b**, Whole mount EdU staining and tubulin expression during CNS regeneration. All panels are anterior to the top, right view. Whole mount staining of CNS regeneration at 16 h, 2, 5, and 7 dpa showing newly divided EdU positive cells in the regenerated tissue and in regenerating nerve fibers expressing tubulin. Enlargements of the square area appear in the following image in the panel. At 16 hpa the wound is open and the neural complex is absent. Cilia in the exposed branchial basket (BB) are positive for tubulin. At 2 dpa the wound is closed but the neural complex is still absent. EdU positive cells can be seen in the regenerated body wall. At 5 and 7 dpa we see the regenerated neural complex (NC) composed of tubulin and EdU positive cells. Anterior siphon (AS) and oral siphon (OS). **c-d**, Quantification of EdU-positive cells in 100µm^2^ sections (n=4 sections per animal, n=3 animals per time point). Violin plots display the number of EdU-positive cells in each section. Data are mean ± SE. *P* values determined by Mann Whitney Wilcoxon test and are indicated above each boxplot. **c**, Number of EdU-positive cells in the regenerating tissue along the different time points (*n* = 3 per time point). **d**, Number of EdU-positive cells in the origin and regenerated tissue along the different time points (*n* = 3 per time point).

**Extended data Fig. 4.**
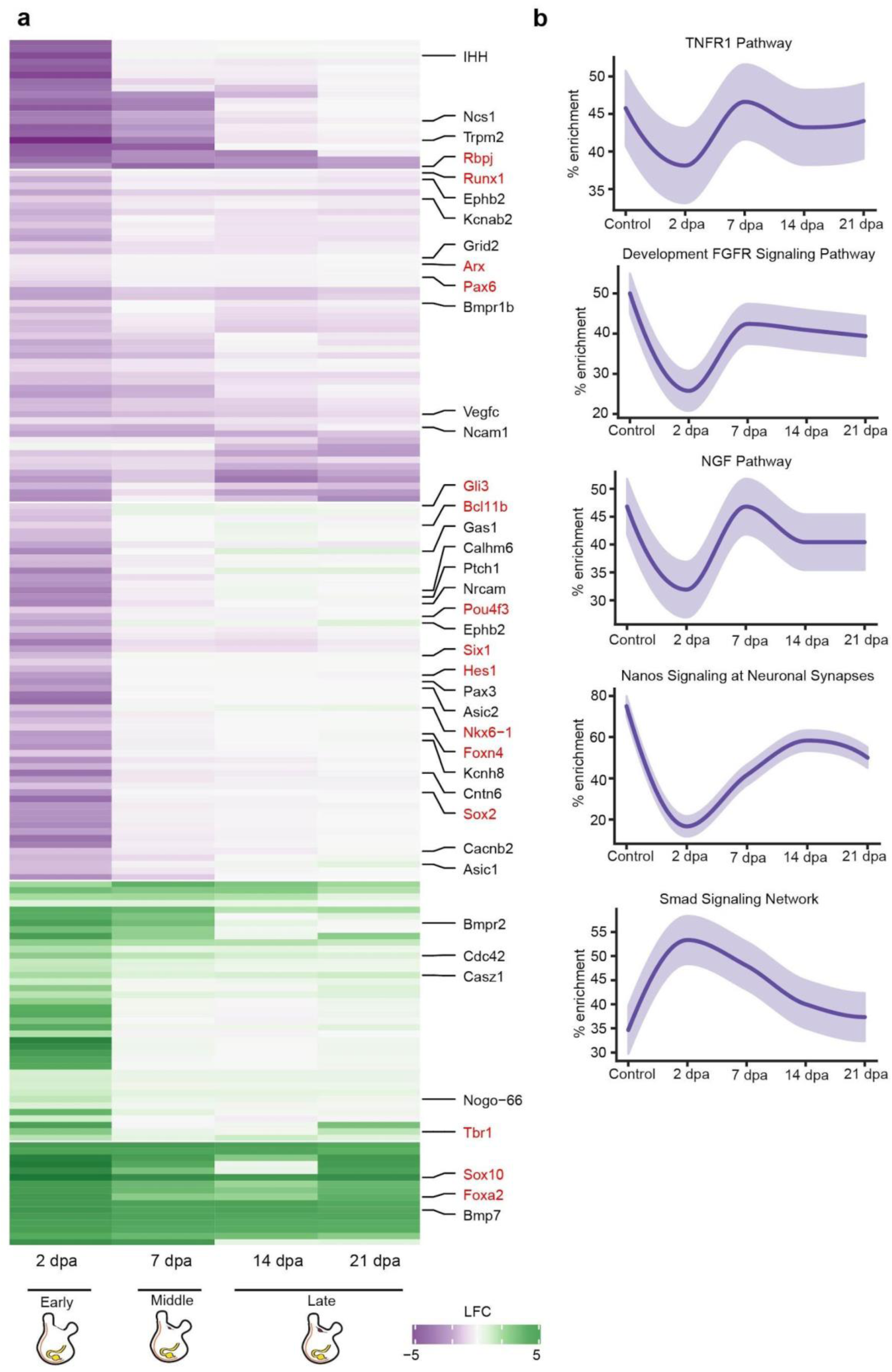
Expression profile for genes involved in nerve regenerative functions. **a**, Heatmap depicts log fold change values for significantly expressed genes (rows) over the sampled regeneration time points. Green indicates a positive fold change (upregulated with respect to uncut CNS), and purple indicates a negative fold change (downregulated with respect to control). Genes of interest appear on the right. Regulatory factors are colored red^82^. **b**, Gene enrichment plot of regeneration-associated genes during CNS regeneration. Light-shaded regions indicate the 50% and 99% confidence intervals under a hypergeometric model.

**Extended data Fig. 5.**
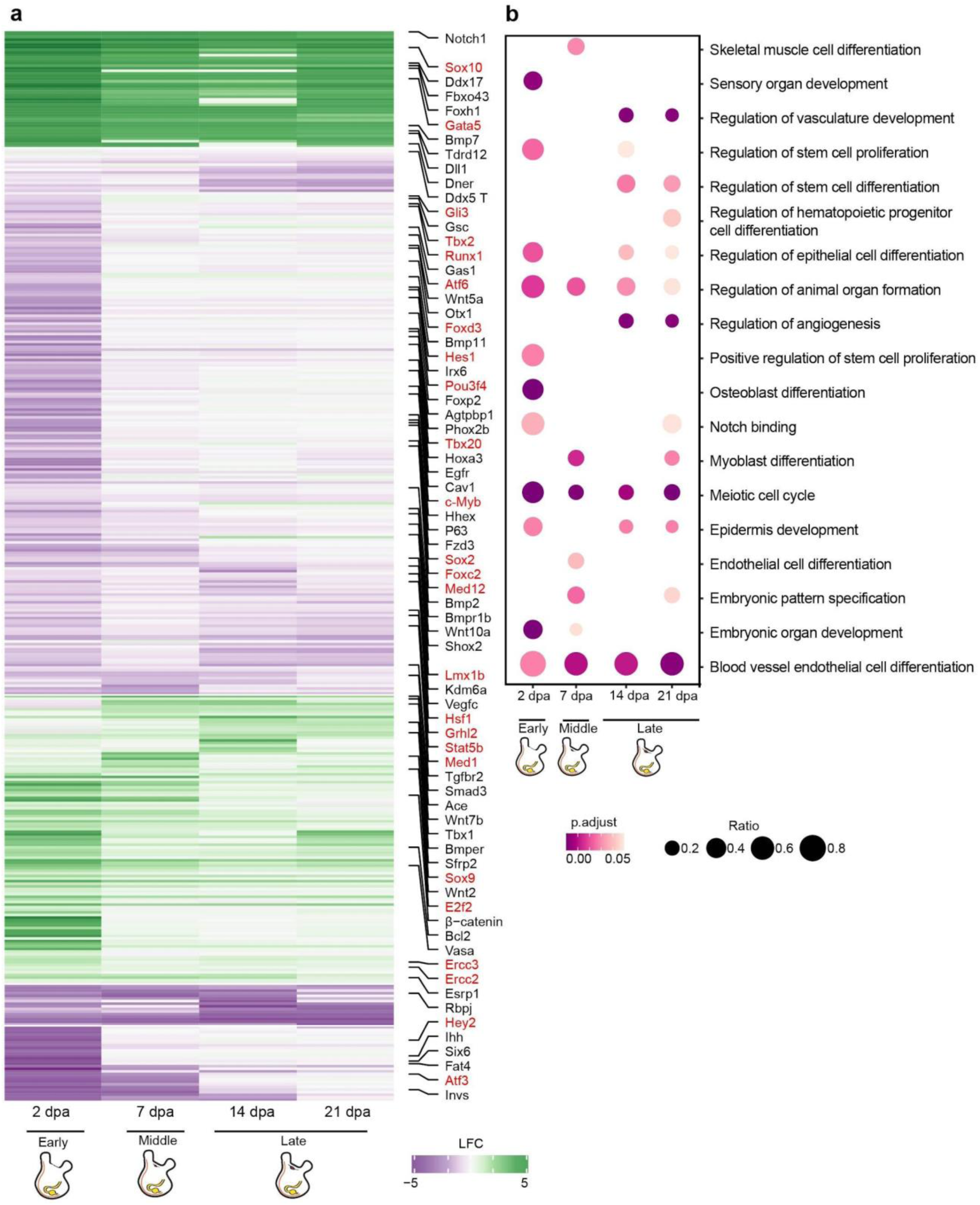
Expression profile for genes involved in regenerative functions. **a,** Heatmap depicts log fold change values for significantly expressed genes (rows) over the sampled regeneration time points. Green indicates a positive fold change (upregulated with respect to uncut CNS), and purple indicates a negative fold change (downregulated with respect to control). Genes of interest appear on the right. Transcription factors are colored red^82^. **b,** Regeneration related gene ontology (GO) term enrichments for each of the time points (ORA, FDR≤0.05).

